# Beyond Mean Pooling: Statistical Meta-Aggregation for Subject-Level EEG Classification in Parkinson’s Disease

**DOI:** 10.64898/2026.10.13.759017

**Authors:** Taghipour L. Pouya, Amirsadra Khodadadi, Faranak Rezaei, Zahra Anvarian, Roberto C. Sotero, Sadegh Madadi

**Affiliations:** University of Calgary, Department of Biomedical Engineering; Hotchkiss Brain Institute; Amirkabir University of Technology, Department of Electrical Engineering; University of Calgary, Department of Computer Science; Amirkabir University of Technology, Department of Biomedical Engineering

**Author notes:** Equal contribution with A. Khodadadi. Equal contribution with P. Taghipour L. Equal contribution with Z. Anvarian. Equal contribution with F. Rezaei.

## Abstract

Electroencephalography (EEG) provides a non-invasive, temporally precise tool for studying neural alterations associated with Parkinson’s disease (PD). When labels are defined at the subject level but inputs are epoch-level, a key design choice is how epoch-wise predictions are aggregated to a single subject-level decision. Most prior work uses mean pooling, which assumes all epochs contribute equally and independently. This study proposes a statistical meta-aggregation strategy that represents each subject by 14 descriptors of the epoch-wise prediction distribution, including measures of central tendency, dispersion, shape, and high-confidence concentration. Using a self-supervised contrastive encoder trained on task-evoked EEG from the Simon conflict task, and a sample of 90 subjects including 57 PD and 33 control, this work evaluates the proposed framework under subject-wise five-fold cross-validation. Replacing mean pooling with statistical meta-aggregation lifts subject-level ROC-AUC from 0.655 to 0.872 for logistic regression and from 0.724 to 0.840 for support vector machines, without modifying the encoder or the classifier family. Averaging predictions across three independent self-supervised pretraining seeds yields competitive performance and improved balanced accuracy. These results suggest that, in this dataset, the way epoch-level predictions are summarized at the subject level has a substantial effect on final PD classification performance.

## 1 Introduction

Parkinson’s disease is a progressive neurodegenerative disorder affecting approximately 1% of the population over the age of 60. In addition to its motor signs, PD is accompanied by cognitive impairments and executive dysfunction that may precede clinical diagnosis by years, motivating the search for objective, non-invasive biomarkers [1, 2]. Electroencephalography is well suited to this role since it is affordable, portable, and offers millisecond temporal resolution, supporting both clinical and home-based implementations [3, 4].

Most automated EEG-based PD classification methods fall into two families: classical pipelines that combine handcrafted spectral, statistical, or higher-order features with linear classifiers [5–7], and deep models that learn spatiotemporal representations end-to-end [8– 11]. A largely separate line of research uses task-based paradigms such as the Simon conflict task to probe specific executive-control mechanisms that are impaired in PD, with mid-frontal theta during conflict and error monitoring as a robust neural signature [12–14].

Across all of these approaches, EEG classification is naturally a hierarchical learning problem. Each subject contributes many short epochs, but the diagnostic label is defined only at the subject level. Most pipelines convert epoch-level model outputs to a subject-level decision through mean pooling, implicitly assuming that every epoch is an equally informative draw from the same underlying distribution. This assumption is not borne out in practice; Task-evoked EEG is highly variable across trials, and the diagnostic content of a single epoch can vary with attention, fatigue, conflict condition, and signal quality. A subject who produces a wide range of confident and unconfident epochs is materially different from a subject who produces a narrow distribution centered on the same mean, yet mean pooling renders these cases identical.

Self-supervised learning (SSL) has emerged as a strong paradigm for EEG representation learning under label scarcity, with contrastive approaches such as SimCLR, MoCo, BYOL, and time-series variants demonstrating utility in sleep staging, emotion recognition, and motor imagery [15–22]. However, SSL applications to task-based EEG and to PD classification remain limited, and the aggregation step downstream of the encoder has received little attention.

This work re-examines the aggregation step itself. This study proposes that the distribution of epoch-wise predictions for a subject is a richer signal than its mean. Each subject is represented by 14 statistical descriptors of that distribution, and a logistic regression or support vector machine is trained on these descriptors. The encoder, the data, the cross-validation splits, and the classifier family are held fixed; only the aggregation function changes. Under these controlled conditions, the lift attributable to meta-aggregation is across both classifier choices. First, this work evaluates whether subject-level EEG classification can be improved by summarizing the full distribution of epoch-wise prediction scores rather than only their mean. Second, using the same encoder, folds, and classifier families, this research compares mean pooling with a 14-feature statistical aggregation scheme. Third, testing whether averaging predictions across three SSL pretraining seeds improves robustness under class imbalance was applied.

This work used the publicly available Simon Conflict Task dataset (OpenNeuro ds004580, v1.0.0) [23], distributed under the CC0 license and organized in the Brain Imaging Data Structure standard [24]. After quality control (subject-level event-alignment checks and channel-availability filtering), the dataset used in this study comprises 90 subjects, where there are 57 with clinically diagnosed Parkinson’s disease and 33 age-matched healthy controls. Continuous scalp EEG was acquired with a BrainVision system at 500 Hz using the extended 10–20 system; participants performed the classical Simon task in which they responded to the colour of a lateralized stimulus while ignoring its spatial location, generating congruent and incongruent trials [12, 25].

## 2 Method

### 2.1 Data Preprocessing

Preprocessing was implemented in MNE-Python [26] via MNE-BIDS [27]. Continuous data were band-pass filtered between 0.1 and 40 Hz (fourth-order zero-phase Butterworth), rereferenced to the common average, and decomposed with Independent Component Analysis (Infomax) after PCA whitening, retaining 99% of variance [28, 29]. Components were classified by ICLabel [30]; components with posterior probability above 0.70 for any non-neural category were removed. Stimulus-locked epochs were extracted from −0.2 to 0.8 s with baseline correction over [−0.2, 0] s. After QC, the 60 EEG channels common to all retained subjects were kept. Each epoch was z-scored per channel across its time samples prior to model input. Subjects retained a mean of 333 epochs (range 115–380), for a total of 29 958 epochs.

### 2.2 Self-Supervised Encoder

The encoder is a compact convolutional network operating on inputs of shape (1, *C, T*) with *C*=60 channels and *T* =500 time points. It consists of a temporal convolution (16 filters, kernel (1, 25)), a spatial convolution (32 filters, kernel (*C*, 1)), and a second temporal convolution (64 filters, kernel (1, 15)), each followed by batch normalization and ELU activation [31]. Average pooling and a linear projection yield a 128-dimensional embedding, which a two-layer MLP head maps to a 64-dimensional contrastive space. The model is trained with NT-Xent loss [16] on EEG-specific augmentations (additive Gaussian noise, temporal masking, amplitude scaling) without using diagnostic labels. To control compute and prevent any single subject from dominating training, the number of training epochs per subject is capped at 250 via random subsampling.

### 2.3 Subject-Level Aggregation Schemes

Given a trained encoder, each epoch is projected to an embedding. A linear classifier (logistic regression or SVM) is trained to produce per-epoch probabilities of PD. The question studied in this paper is how these per-epoch probabilities are aggregated to a single subject-level decision. We compare two schemes, which are

#### Mean pooling (baseline)

For subject *s* with epoch probabilities 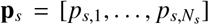 the subject-level score is 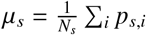 and the label is 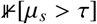 for threshold τ.

##### Statistical meta-aggregation (proposed)

The subject is represented by a vector of 14 distributional descriptors:

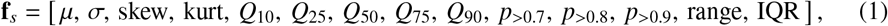

covering central tendency (µ, *Q*_50_), dispersion (σ, range, IQR), distribution shape (skewness, kurtosis, percentiles), and the proportion of high-confidence epochs at three thresholds. The 14 features are z-scored across subjects within each training fold and used as input to a logistic regression or RBF-kernel SVM classifier. Subject-level labels are obtained by thresholding the classifier’s probability output at τ = 0.4, the value used throughout this study.

### 2.4 Cross-Validation and Ensembling

All evaluations use subject-wise five-fold cross-validation, where every epoch from a given subject appears in exactly one partition per fold, so no subject’s data leaks across train and test. The self-supervised encoder is pretrained independently for three random seeds (42, 123, 999); embeddings from each seed feed the downstream aggregation and classification stages. We also evaluate a *meta-ensemble*, in which the subject-level PD probabilities from the three seed-specific meta-aggregation models are averaged before thresholding.

## 3 Results

All metrics are reported at the subject level, in line with the clinical setting where decisions are made per patient rather than per trial. This study reports accuracy, balanced accuracy, and ROC-AUC, and presents both the fold-mean ROC-AUC (mean of per-fold AUCs across the five subject-wise folds) and the pooled ROC-AUC (computed by concatenating subject-level predicted probabilities across folds and computing one AUC). Both summaries are legitimate; fold-mean is the primary metric because it reflects between-fold variability, while pooled AUC corresponds to the single curve plotted in the ROC figures.

### 3.1 Meta-Aggregation Substantially Outperforms Mean Pooling

Table 1 reports subject-level performance for the mean-pooling baseline (linear classifiers on mean-pooled SSL embeddings) and the proposed meta-aggregation scheme.

**Table 1.** Subject-level performance: mean pooling vs. statistical meta-aggregation. Values are means across five subject-wise folds; for the mean-pooling baseline, the standard deviation across folds is also reported. Both methods use the same self-supervised encoder and the same subject-wise five-fold cross-validation.

| Method | Accuracy | Balanced Acc. | ROC-AUC |
| --- | --- | --- | --- |
| SSL + mean pooling + LR | 0.633 ± 0.075 | 0.607 ± 0.070 | 0.655 ± 0.178 |
| SSL + mean pooling + SVM | 0.644 ± 0.067 | 0.620 ± 0.078 | 0.724 ± 0.068 |
| SSL + meta-aggregation + LR | 0.722 | 0.730 | **0.872** |
| SSL + meta-aggregation + SVM | 0.644 | 0.650 | 0.840 |

Replacing mean pooling with statistical meta-aggregation lifts the ROC-AUC by 0.217 for logistic regression (from 0.655 to 0.872) and by 0.116 for the SVM (from 0.724 to 0.840). Balanced accuracy improves by 0.123 and 0.030, respectively. These gains are obtained without modifying the encoder, the classifier family, the data, or the cross-validation splits; only the aggregation function changes. Because the encoder, data, classifier family, and subject-wise folds were kept fixed, the improvement is most likely explained by the aggregation stage rather than by a change in representation learning.

### 3.2 Per-Fold Stability and Fold 3

Table 2 reports per-fold metrics for both meta-aggregation classifiers.

**Table 2.** Per-fold subject-level metrics for the proposed meta-aggregation pipeline.

| Fold | Meta-Aggregation + LR |  |  | Meta-Aggregation + SVM |  |  |
| --- | --- | --- | --- | --- | --- | --- |
|  | Acc. | Bal. Acc. | AUC | Acc. | Bal. Acc. | AUC |
| 1 | 0.722 | 0.746 | 0.892 | 0.667 | 0.708 | 0.877 |
| 2 | 0.722 | 0.722 | 0.938 | 0.500 | 0.500 | 0.988 |
| 3 | 0.667 | 0.675 | 0.725 | 0.667 | 0.675 | 0.812 |
| 4 | 0.667 | 0.696 | 0.893 | 0.667 | 0.696 | 0.732 |
| 5 | 0.833 | 0.812 | 0.909 | 0.722 | 0.669 | 0.792 |
| Mean | 0.722 | 0.730 | 0.872 | 0.644 | 0.650 | 0.840 |

Fold 3 is the weakest for both classifiers, with LR dropping to AUC 0.725 and SVM to 0.812. The overall pattern is consistent, where meta-aggregation with logistic regression yields AUC above 0.89 in four of five folds. Figure 1 shows the corresponding fold-wise trajectory. The SVM exhibits a wider per-fold range, with one fold (Fold 2) producing a perfect ranking (AUC 0.988) but collapsed thresholded balanced accuracy (0.500). This pattern, a high AUC with poor accuracy, indicates a calibration rather than separation issue and motivates the use of AUC as the primary metric for evaluating the proposed aggregation scheme.

**Figure 1.**
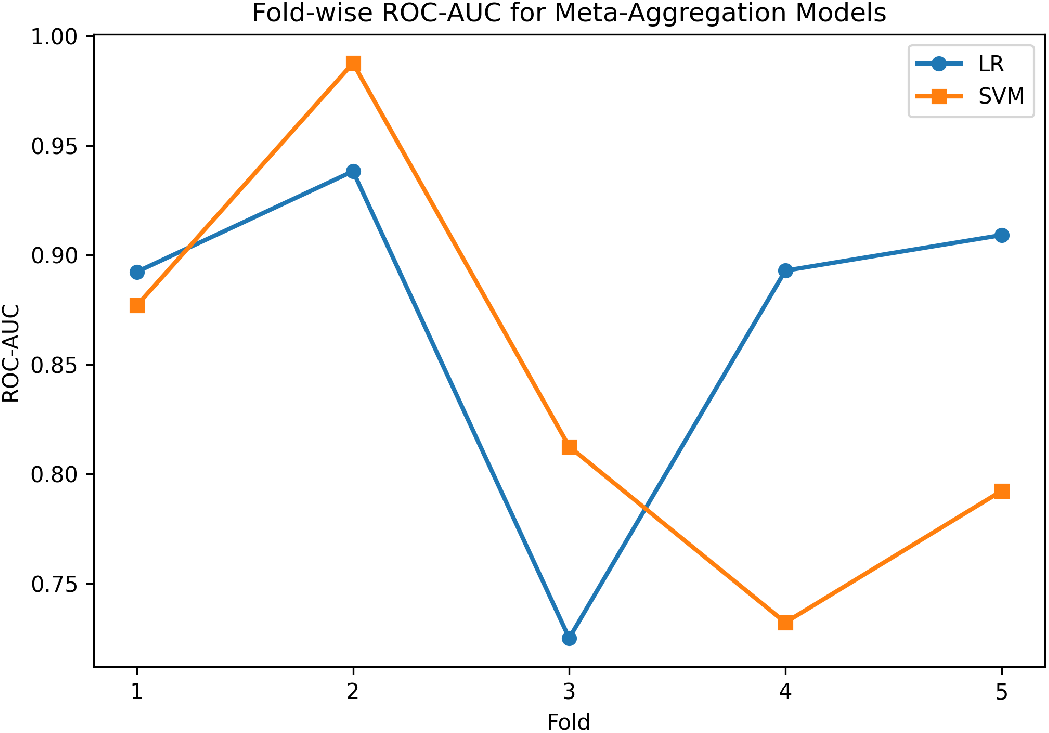
Per-fold ROC-AUC for the two meta-aggregation classifiers. Logistic regression is more stable across folds, while the SVM shows a wider range driven by fold-level threshold calibration.

### 3.3 ROC Curves and Pooled Performance

Figure 2 shows mean ROC curves with shaded between-fold variability for the three meta-aggregation variants. Figure 3 shows the corresponding pooled ROC curves, computed by concatenating subject-level predictions across folds. Pooled AUC values are LR 0.809, SVM 0.746, and Meta-Ensemble 0.791; the gap between fold-mean and pooled AUC reflects the heterogeneity across folds, particularly the weaker Fold 3.

**Figure 2.**
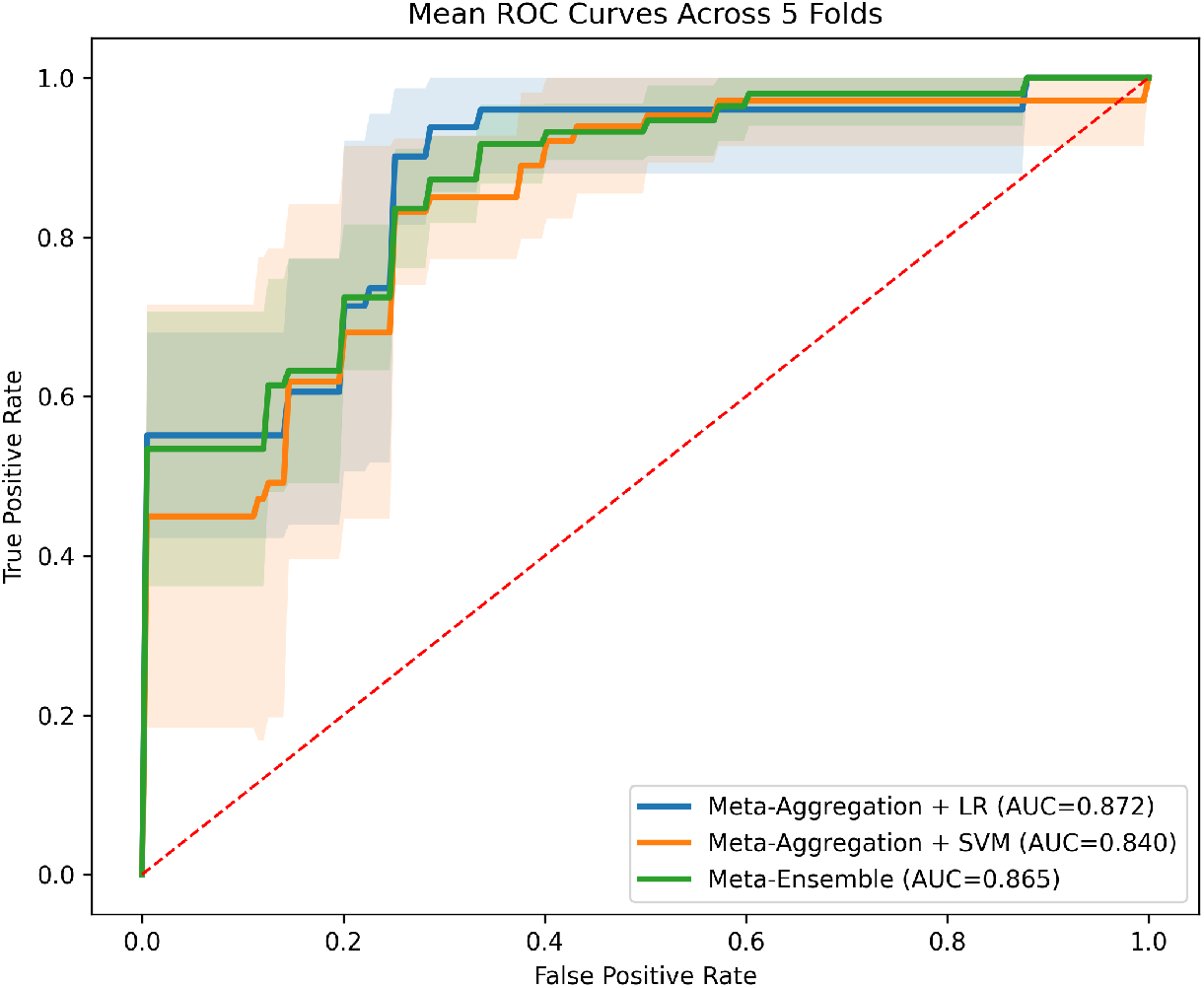
Mean ROC curves across five subject-wise folds. The LR variant gives the highest fold-mean AUC, while shading shows between-fold variability.

**Figure 3.**
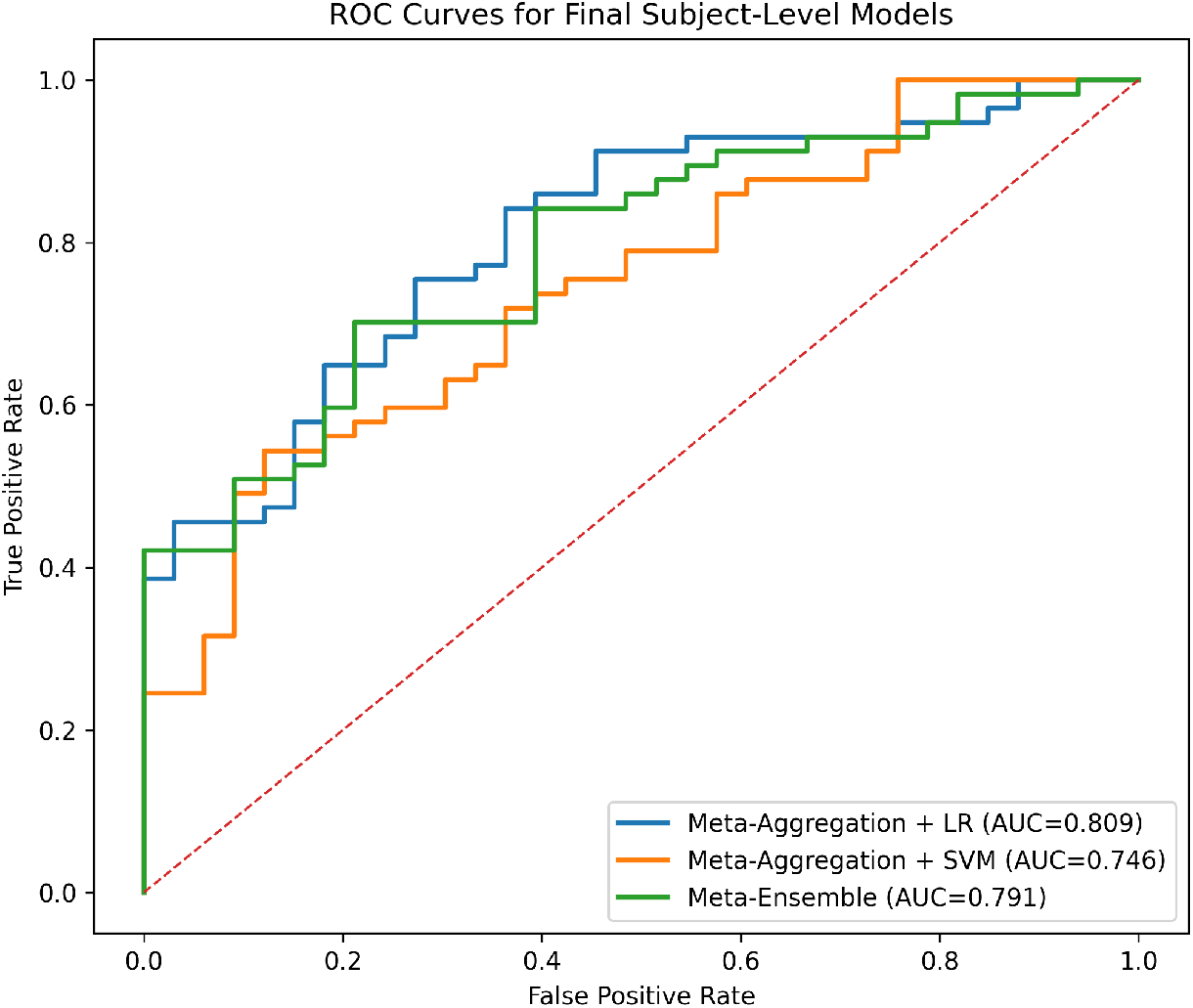
Pooled ROC curves from concatenated subject-level predictions across folds.

### 3.4 Subject-Level Operating Point

At the operating threshold τ = 0.4, the proposed Meta-Aggregation + LR model produces the confusion matrix shown in Figure 4: 43 of 57 PD subjects are correctly identified (sensitivity 0.754) while 22 of 33 controls are correctly identified (specificity 0.667), yielding subject-level accuracy 0.722 and balanced accuracy 0.711. The model recovers most PD subjects without degenerating to the majority class.

**Figure 4.**
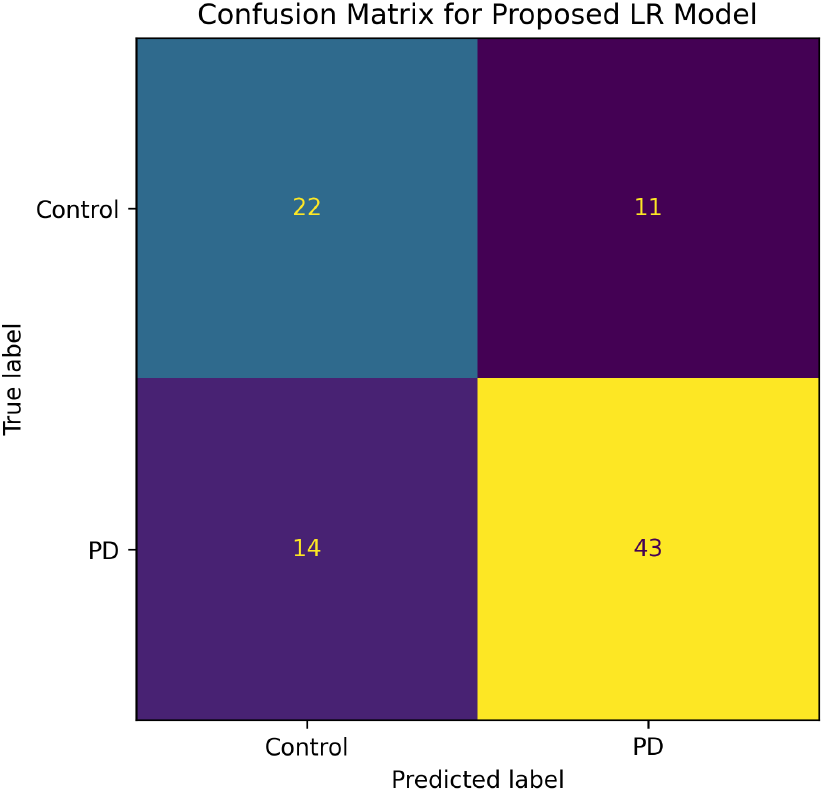
Subject-level confusion matrix for the proposed Meta-Aggregation + LR pipeline at threshold τ = 0.4, pooled across the five test folds (total 90 subjects).

### 3.5 Ensembling Across Pretraining Seeds

Table 3 summarizes the three-seed meta-ensemble.

**Table 3.** Across SSL seeds, averaging predictions improves balanced accuracy and pooled ROC-AUC over individual seeds, suggesting better robustness to pretraining variability.

| Model | Accuracy | Bal. Acc. | ROC-AUC (pooled) |
| --- | --- | --- | --- |
| Seed 42 | 0.700 | 0.624 | 0.720 |
| Seed 123 | 0.722 | 0.676 | 0.764 |
| Seed 999 | 0.633 | 0.573 | 0.695 |
| Meta-Ensemble (avg. of 3 seeds) | 0.711 | **0.727** | 0.791 |

Averaging predictions across seeds improves balanced accuracy from a per-seed maximum of 0.676 to 0.727, and pooled ROC-AUC from 0.764 to 0.791. The ensemble is competitive with the single-seed Meta-Aggregation + LR pipeline (fold-mean AUC 0.865 in the mean-ROC curve) and provides additional robustness against per-seed variability without further model complexity.

### 3.6 The Distribution Carries the Signal

To illustrate why distributional descriptors outperform the mean, Figure 5 plots epoch-level prediction distributions for three example subjects. Mean pooling would summarize a subject with spread-out predictions by a value around 0.5 and is unable to express the difference between this case and a subject with all epochs near 0.5. The proposed descriptors capture this difference directly through σ, the interquartile range, and the high-confidence proportions.

**Figure 5.**
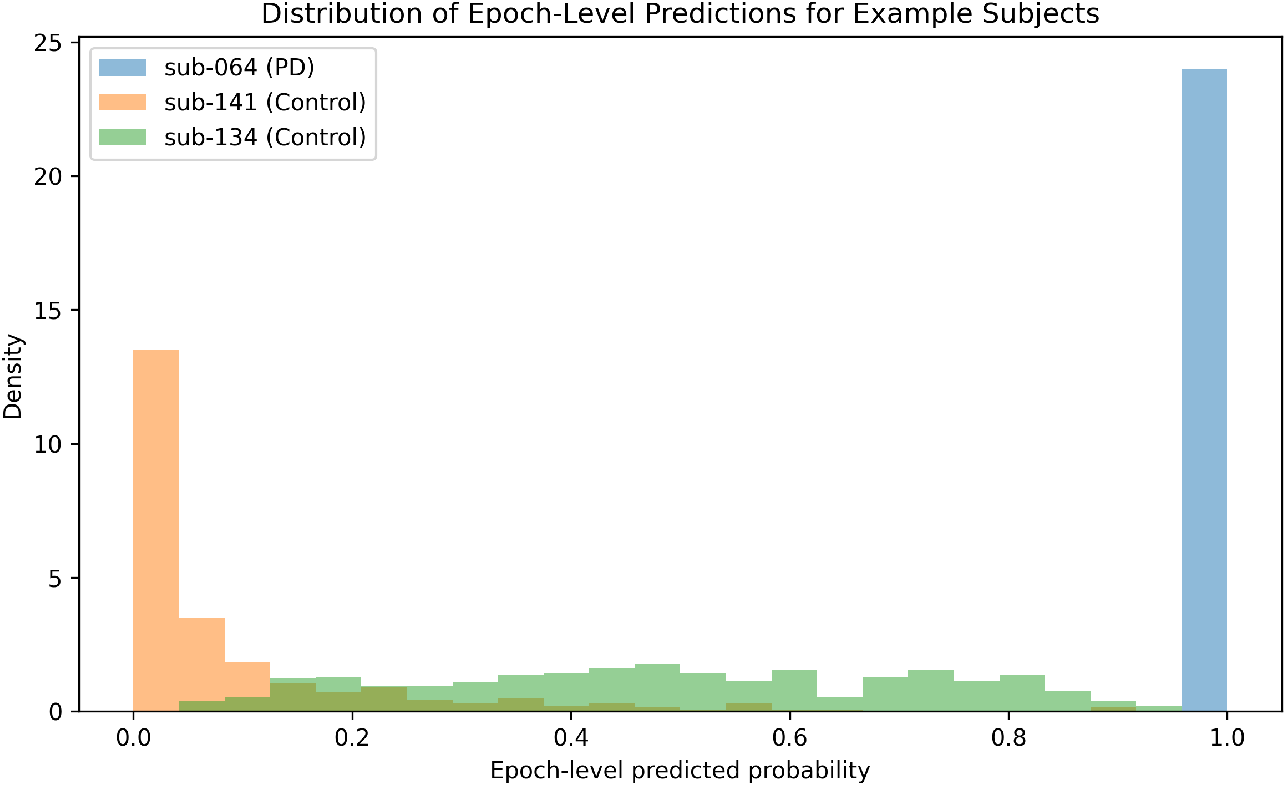
Epoch-level prediction distributions for three example subjects illustrating distinct distribution shapes that are conflated by mean pooling but preserved by the proposed statistical descriptors.

## 4 Conclusion

This study proposed a statistical meta-aggregation approach for subject-level EEG classification in Parkinson’s disease. Instead of reducing all epoch-wise predictions to a single mean score, each subject was represented using 14 descriptors of the prediction-score distribution. On the Simon-task EEG dataset, this change improved subject-level ROC-AUC for both logistic regression and SVM when the encoder, classifier family, data, and subject-wise cross-validation folds were held fixed. The improvement was largest for logistic regression, where ROC-AUC increased from 0.655 with mean pooling to 0.872 with meta-aggregation. These results suggest that, for hierarchical EEG datasets, subject-level aggregation can meaningfully affect performance and should be treated as part of the model design rather than as a minor post-processing step.

Several limitations should be considered. First, the proposed aggregation treats epochs as exchangeable samples and discards their temporal order; fatigue, learning effects, and trial-condition contrasts (congruent vs. incongruent) are not modelled and may carry additional diagnostic information [32]. Future work could replace fixed descriptors with attention-based or sequence-based aggregation that learns which epochs are most informative [33]. Second, the dataset has more PD subjects than controls (57 vs. 33), and although we report balanced accuracy and use subject-wise cross-validation, more sophisticated imbalance treatments could be explored [34]. Third, only three self-supervised pretraining seeds are used; the variability across seeds (per-seed AUC 0.695–0.764) suggests that a larger ensemble may further stabilize performance. Fourth, the results are obtained on a single public dataset and a single task; transfer to other paradigms, sites, and recording systems remains to be established [35]. Fifth, the framework is not yet directly compared against fully supervised deep models or against classical handcrafted-feature baselines on the same subject set; such comparisons are the natural next step. Finally, the proposed descriptors are not designed to identify which neural sources or frequency bands drive the decision; mechanistic interpretability is left to future work.

The pipeline is computationally light, interpretable at the subject level, and agnostic to the choice of representation. Beyond Parkinson’s disease, the same principle applies wherever EEG decisions must be aggregated from many short epochs to a single subject-level outcome, including sleep staging, anaesthesia monitoring, and seizure risk stratification. We release the subject-level prediction files and aggregation code to support replication.

